# CRISPR-Cas9 human gene replacement and phenomic characterization in *Caenorhabditis elegans* to understand the functional conservation of human genes and decipher variants of uncertain significance

**DOI:** 10.1101/369249

**Authors:** Troy A. McDiarmid, Vinci Au, Aaron D. Loewen, Joseph Liang, Kota Mizumoto, Donald G. Moerman, Catharine H. Rankin

**Author notes:** Correspondence concerning this article should be addressed to: Catharine H. Rankin, Djavad Mowafaghian Center for Brain Health and Department of Psychology, University of British Columbia, 2136 West Mall, Vancouver, British Columbia, Canada, V6T 1Z4.

## Abstract

Our ability to sequence genomes has vastly surpassed our ability to interpret the genetic variation we discover. This presents a major challenge in the clinical setting, where the recent application of whole exome and whole genome sequencing has uncovered thousands of genetic variants of uncertain significance. Here, we present a strategy for targeted human gene replacement and phenomic characterization based on CRISPR-Cas9 genome engineering in the genetic model organism *Caenorhabditis elegans* that will facilitate assessment of the functional conservation of human genes and structure-function analysis of disease-associated variants with unprecedented precision. We validate our strategy by demonstrating that direct single-copy replacement of the *C. elegans* ortholog (*daf-18*) with the critical human disease-associated gene Phosphatase and Tensin Homolog (*PTEN*) is sufficient to rescue multiple phenotypic abnormalities caused by complete deletion of *daf-18*, including complex chemosensory and mechanosenory impairments. In addition, we used our strategy to generate animals harboring a single copy of the known pathogenic lipid phosphatase inactive PTEN variant (PTEN-G129E) and showed that our automated *in vivo* phenotypic assays could accurately and efficiently classify this missense variant as loss-of-function. The integrated nature of the human transgenes allows for analysis of both homozygous and heterozygous variants and greatly facilitates high-throughput precision medicine drug screens. By combining genome engineering with rapid and automated phenotypic characterization, our strategy streamlines identification of novel conserved gene functions in complex sensory and learning phenotypes that can be used as *in vivo* functional assays to decipher variants of uncertain significance.

## Introduction

The rapid development and application of whole exome and whole genome sequencing technology has dramatically increased the pace at which we associate genetic variation with a particular disease (Auton et al., 2015; Bamshad et al., 2011; Gonzaga-Jauregui et al., 2012; Lek et al., 2016; Metzker, 2010; Need et al., 2012; Ng et al., 2010). However, our ability to sequence genomes has vastly surpassed our ability to interpret the clinical implications of the genetic variants we discover. The majority of genetic variants identified in clinical populations are currently classified as “variants of uncertain significance” meaning their potential role as a causative agent in the disease in question, or their pathogenicity, is unknown (Richards et al., 2015). Many variants are exceedingly rare, making it extremely difficult to designate them as pathogenic using classical genetic methods such as segregation within a pedigree or by identifying multiple carriers of the variant. As such, it often remains challenging to predict clinical outcomes and make informed treatment decisions based on genetic data alone.

In an attempt to address this problem several computational tools have been developed that estimate the functional consequences and pathogenicity of disease-associated variants (Richards et al., 2015). These tools use a variety of predictive features such as evolutionary sequence conservation, protein structural and functional information, the prevalence of a variant in large putatively healthy control populations, or a combination of annotations (Kircher et al., 2014; Lek et al., 2016; Richards et al., 2015). Despite extensive efforts none of these tools used in isolation or combination can faithfully report on the functional effects of a large portion of disease-associated variation and their accuracy is intrinsically limited to existing experimental training data (Grimm et al., 2015; Miosge et al., 2015; Starita et al., 2017). These limitations were clearly demonstrated in a recent study that showed *in vivo* functional assays of 21 human genes in yeast identified pathogenic variants with significantly higher precision and specificity than computational methods (Sun et al., 2016). This means that even for genes with well-characterized biological functions there are often hundreds of variants of uncertain functional significance (Landrum et al., 2014; Starita et al., 2017). This creates a challenging situation that requires direct assessment of the functional effects of disease-associated variants *in vivo* (Starita et al., 2017).

Genetically tractable model organisms are critical for discovering novel gene functions and the functional consequences of disease-associated genetic variants (Dunham and Fowler, 2013; Lehner, 2013; Manolio et al., 2017; Wangler et al., 2017). Governmental and private funding agencies are increasingly commissioning large-scale collaborative programs to use diverse genetic model organisms to decipher variants of uncertain significance (Chong et al., 2015; Gahl et al., 2016; Wangler et al., 2017). Among genetic model organisms, the nematode *Caenorhabditis elegans* has proven to be a particularly powerful animal model for the functional characterization of human genes *in vivo* (Kaletta and Hengartner, 2006). *C. elegans* is an ideal genetic model as it combines the throughput and tractability of a single-celled organism with the complex morphology and behavioral repertoire of a multicellular animal. In addition, approximately 60-80% of human genes have an ortholog in the *C. elegans* genome (Kaletta and Hengartner, 2006; Lai et al., 2000a; Shaye and Greenwald, 2011). Transgenic expression of human genes is routinely done to confirm functional conservation and to observe the effects of disease-associated mutations. Notable examples of the utility of *C. elegans* to determine conserved human gene functions relevant to disease include the identification of presenilins as part of the gamma secretase complex, the mechanism of action of selective serotonin reuptake inhibitors, and the role of the insulin signaling pathway in normal and pathological ageing (Kaletta and Hengartner, 2006; Levitan et al., 1996; Levitan and Greenwald, 1995; Murphy et al., 2003; Ranganathan et al., 2001). However, traditional methods for expression of human genes in *C. elegans* rely on mosaic and variable over-expression of transgenes harbored as extrachromosomal arrays or specialized genetic backgrounds that can confound phenotypic analysis. This presents several challenges that inhibit precise analysis of the often critical but subtle effects of missense variants and impede the use of these transgenic strains in large-scale drug screens.

The recent advent of CRISPR-based genome editing has revolutionized structure-function analyses across model organisms (Cong et al., 2013; DiCarlo et al., 2013; Dickinson et al., 2013; Doudna and Charpentier, 2014; Friedland et al., 2013; Gratz et al., 2013; Hwang et al., 2013; Jinek et al., 2013, 2012; Li et al., 2013). This system uses a single guide RNA (sgRNA) to precisely target a nuclease (most often Cas9) to induce a DNA double strand break at a defined location (Doudna and Charpentier, 2014). The double strand break can then be repaired via the error-prone non-homologous end-joining pathway (often resulting in damaging frameshift mutations) or a more precise homology repair pathway, e.g. homology directed repair (HDR) or microhomology-mediated end joining. Following a double strand break, exogenous DNA repair templates can be used as substrates for homologous recombination, allowing virtually any desired sequence to be inserted anywhere in the genome. Importantly, CRISPR-Cas9 genome engineering is remarkably efficient and robust in *C. elegans* (Dickinson and Goldstein, 2016; Norris et al., 2015).

Here, we present a broadly applicable strategy that adapts CRISPR-Cas9 genome engineering for targeted replacement of *C. elegans* genes with human genes. We illustrate how the library of knockout and humanized transgenics generated with this approach can be efficiently combined with automated machine vision phenotyping to rapidly discover novel gene functions, assess the functional conservation of human genes, and how this will allow for analysis of the effects of variants of uncertain significance with unprecedented precision. It is our hope that the human gene replacement and phenomic characterization strategy delineated in this article will serve both basic and health researchers alike, by serving as an open and shareable resource that will aid any genome engineer interested in understanding the functional conservation of human genes, and the functional consequences of their variants.

## Results

### A general genome editing strategy for direct replacement of a *C. elegans* gene with a single copy of its human ortholog

To replace the Open Reading Frame (ORF) of an orthologous gene with a human gene our strategy first directs an sgRNA to induce a Cas9 mediated DNA double strand break immediately downstream of the ortholog start codon (Fig. 1A). A co-injected repair template containing ~500 bp homology arms targeted to the regions immediately upstream and downstream of the ortholog ORF serve as a substrate for homology directed repair. By fusing the coding DNA sequence (CDS) of a human gene of interest to the upstream homology arm homology directed repair integrates the human gene in place of the ortholog at a single copy in frame (Fig. 1A).

**Figure 1.**
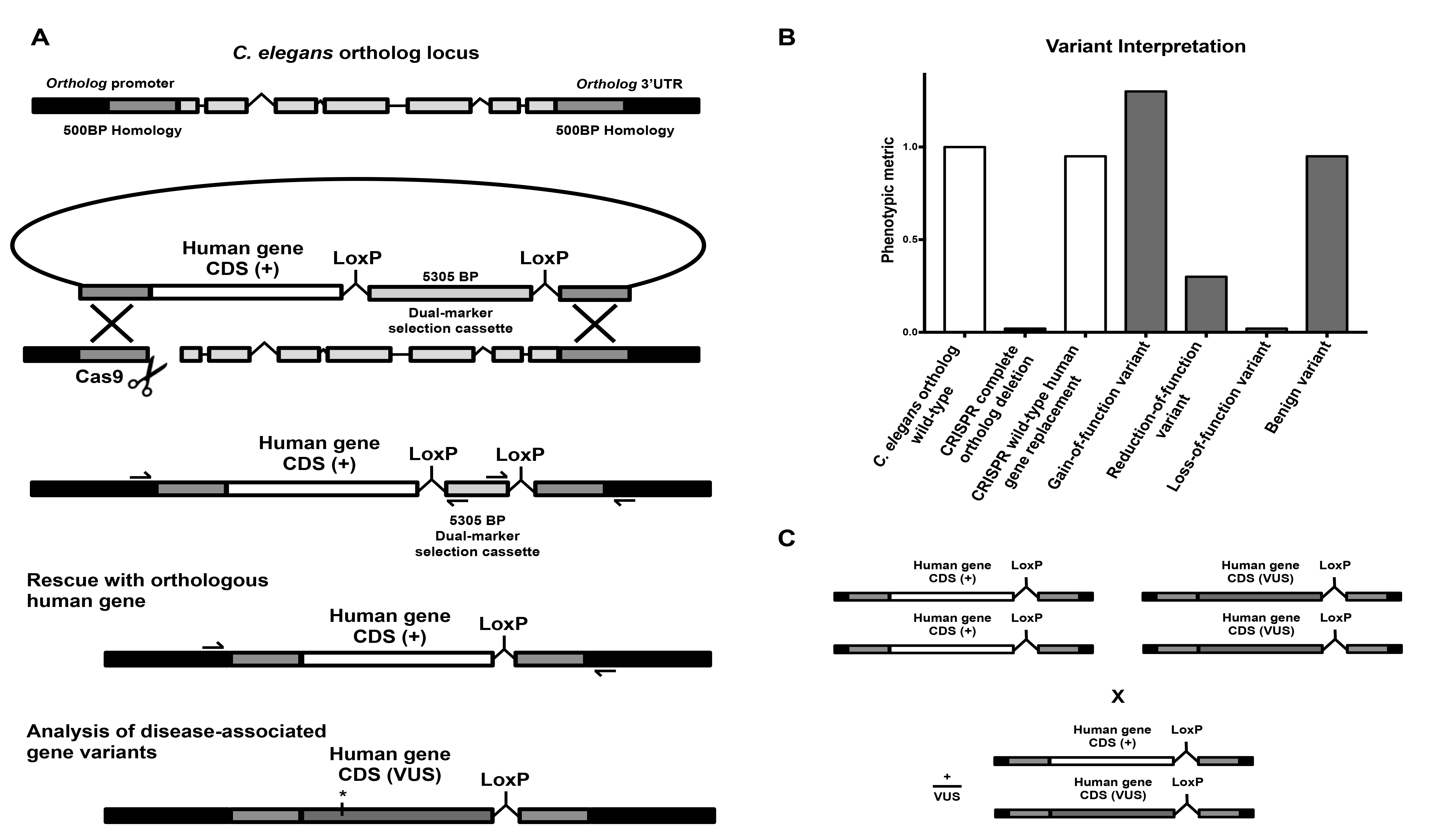
A general strategy for direct single copy replacement of C. elegans genes with human genes at the orthologs native genomic loci. **A)** A schematic of the genome editing strategy. (top left) A sgRNA targets Cas9 to induce a DNA double strand break immediately downstream of the orthologs start codon. A co-injected repair template containing ~500BP homology arms targeted to the regions immediately upstream and downstream of the ortholog ORF serve as a substrate for homology directed repair. By fusing the CDS of a human gene of interest to the upstream homology arm homology directed repair integrates the human gene in place of the ortholog at a single copy in frame. A co-integrated Dual-Marker selection cassette consisting of an antibiotic resistance gene (*Prps-27::neoR::unc-54 UTR*) and a fluorescent marker (*Pmyo-2::GFP::unc-54 UTR*) greatly facilitates the identification transgenic animals without inducing morphological or phenotypic abnormalities. (middle left) initial integration deletes the entire open reading frame of the C. elegans ortholog while separating the human gene from the orthologs transcriptional terminator to inhibit expression, creating an ortholog deletion allele for phenotypic analysis (Note: cassette is not shown to scale for most human gene CDS). (bottom left) Subsequent injection of Cre Recombinase excises the selection cassette and connects the human gene to the orthologous transcriptional termination sequence such that a single copy of the human gene will now be expressed under the control of all of the orthologs 5’ and 3’ cis-and trans-regulatory machinery. Validation of the desired edit is preformed using standard amplification and Sanger sequencing of the target region (primer binding locations represented by half arrows in the schematic). For analysis of a human gene variant of uncertain significance (VUS) the variant of interest is incorporated into the HDR plasmid using standard *in vitro* methods such as site-directed mutagenesis and the same genome editing process is repeated using the same validated sgRNA and homology arms. **B)** Human gene replacement allows for straight-forward interpretation of variant functional effect. This process allows for: 1) initial generation and phenotypic analysis of a complete null allele in the *C. elegans* orthologous gene 2) Direct integration of the human gene to determine if the human gene can compensate for loss of the orthologous gene, measuring functional conservation, 3) structure-function analysis of the effects of variants of uncertain significance on WT gene function. **C)** This strategy allows for straightforward assessment of heterozygous alleles using standard genetic crosses.

To streamline genome editing we have based our method on a recently described Dual-Marker Selection (DMS) cassette screening protocol (Norris et al., 2015). The DMS Cassette consists of an antibiotic resistance gene (*Prps-27::neoR::unc-54 UTR*) and a fluorescent marker (*Pmyo-2::GFP::unc-54 UTR*) that greatly facilitates the identification transgenic animals (Norris et al., 2015). We chose this cassette over similar methods as it: 1) can be used in any wild-type or mutant strain amenable to transgenesis and does not require any specialized genetic backgrounds, and 2) avoids the use of morphology and/or behavior altering selection markers that necessitate cassette excision prior to phenotypic analysis (Bend et al., 2016; Dickinson et al., 2015, 2013). In our strategy, the DMS cassette is placed between the human gene of interest and the downstream homology arm (Fig. 1A). This deletes the entire ORF of the ortholog and separates the human gene from the orthologs transcriptional terminator upon initial integration. In many cases this efficiently creates a useful deletion allele of the ortholog with no human gene expression. In the unlikely event human gene expression does occur without a transcription terminator a second repair template using the same validated homology arms and sgRNA and no human gene can be integrated to create an ortholog null. Importantly, the DMS cassette is flanked by two LoxP sites housed within synthetic introns that allow subsequent excision of the selection cassette via transient expression of Cre Recombinase (Norris et al., 2015). DMS cassette excision connects the human gene to the endogenous *C. elegans* orthologs transcriptional termination sequence such that a single copy of the human gene will now be expressed under the control of all of the orthologs 5’ and 3’ cis-and trans-regulatory machinery (Fig. 1B). Validation of the desired edit is performed using standard PCR amplification and Sanger sequencing of both the 5’ and 3’ junctions of the target region (Fig. 1A).

For structure-function analysis of a human gene variant of uncertain significance the variant of interest is incorporated into the HDR plasmid using standard *in vitro* methods such as site-directed mutagenesis and the same genome editing process is repeated using the same validated sgRNA and homology arms. This process allows for: 1) initial generation and phenotypic analysis of a complete deletion allele in the *C. elegans* orthologous gene, 2) direct integration of the human gene to determine if the human gene can compensate for loss of the orthologous gene, measuring functional conservation, and 3) structure-function analysis of the effects of variants of uncertain significance on wild-type gene function (Fig. 1B). Importantly, the vast majority of variants of uncertain significance identified in patients are heterozygous and the integrated nature of the transgenes generated with this strategy allow for straightforward assessment of heterozygous alleles using standard genetic crosses (Fig. 1D).

### *PTEN* as a prototypic disease-associated gene for targeted human gene replacement

As a proof of principle, we demonstrated the utility of our strategy by focusing on the critical disease-associated gene phosphatase and tensin homolog (*PTEN*). *PTEN* is a lipid phosphatase that antagonizes the phosphoinositide 3-kinase (PI3K) signaling pathway by dephosphorylating phosphatidylinositol (3,4,5)-trisphosphate (PIP3) (Li et al., 1997; Maehama and Dixon, 1998). Heterozygous germline *PTEN* mutations are associated with diverse clinical outcomes including several tumor predisposition phenotypes (collectively called *PTEN* harmatoma tumor syndrome), intellectual disability, and Autism Spectrum Disorders (Hobert et al., 2014; Li et al., 1997; Danny Liaw et al., 1997; McBride et al., 2010; O’Roak et al., 2012; Orrico et al., 2009; Sanders et al., 2015; Varga et al., 2009). Despite extensive study, it is currently impossible to predict the clinical outcome of a *PTEN* mutation carrier using sequence data alone (Matreyek et al., 2018; Mighell et al., 2018). *PTEN* also has several technical advantages that make it an ideal test case: 1) *PTEN* functions in the highly conserved insulin signaling pathway that is well characterized in *C. elegans* (Ozes et al. 2001; Ogg & Ruvkun 1998; Mihaylova et al. 1999, and Fig. 2B). 2) *C. elegans* has a single *PTEN* ortholog called *daf-18* (Fig. 2 A-D) and transgenic overexpression of human *PTEN* using extrachromosomal arrays has been shown to rescue reduced longevity and dauer defective phenotypes induced by mutations in *daf-18* (Liu and Chin-Sang, 2015; Solari et al., 2005). 3) *C. elegans* harboring homozygous *daf-18* null alleles are viable and display superficially normal morphology and spontaneous locomoter behavior (Mihaylova et al. 1999 and Fig. S1).

**Figure 2.**
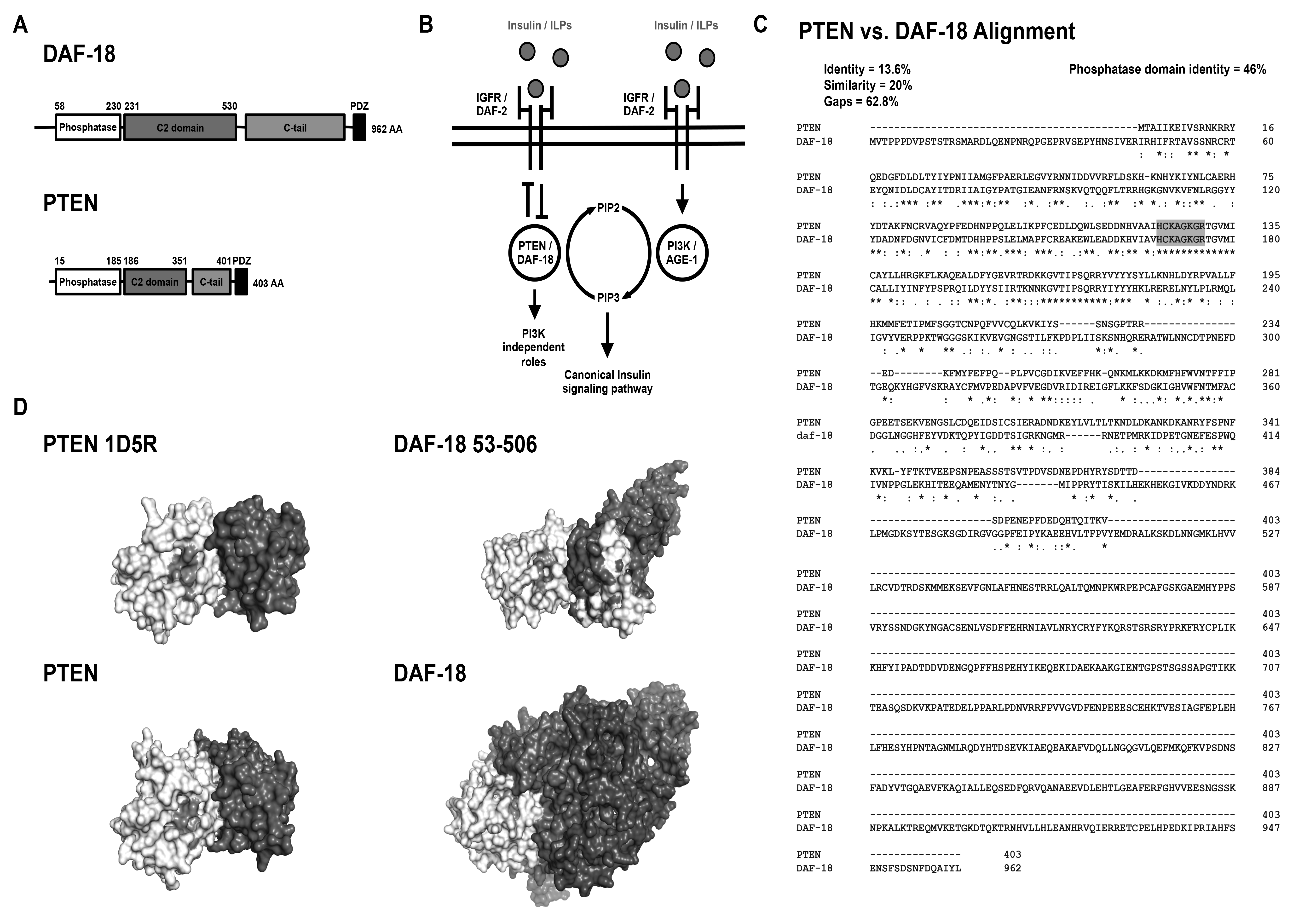
Functional and structural similarity of *C. elegans* DAF-18 and human PTEN. **A)** Protein domain annotations for DAF-18 and PTEN. The canonical DAF-18 amino acid (AA) sequence is more than twice as long as the canonical PTEN AA sequence primarily due to elongation of the C2 membrane targeting and C-tail domains. **B)** Both DAF-18 and PTEN function as lipid and protein phosphatases to antagonize the highly conserved canonical insulin signaling pathway. **C)** Clustal alignment of DAF-18 and PTEN. DAF-18 and PTEN share a highly conserved phosphatase domain (46% identity) and fully conserved catalytic site (residues highlighted in grey). DAF-18 has a markedly longer and less conserved C-terminal region than human PTEN resulting in low overall amino acid similarity (20%) and identity (13.6%). Note that although the C-terminal region is much longer there are small conserved motifs spread throughout that are not illustrated by this alignment (see Liu et al., 2014 oncogene). D) Comparison of DAF-18 and PTEN structural models. **D)** (top left) Solved crystal structure of human PTEN (1D5R reference structure;Lee et al. 1999). (top right) Predicted structure of DAF-18 AA 53-506 indicating similar 3D structure to human PTEN. (bottom left) Predicted structure of full-length PTEN and (bottom right) DAF-18 illustrating the increased size of DAF-18. Note the full length DAF-18 model is likely to be inaccurate due to poor homology-based modeling of the non-conserved C-terminal region. Domain colour mapping matches panel 2A except that the catalytic site within the phosphatase domain is shaded light gray.

### An automated chemotaxis paradigm reveals a conserved nervous system role for PTEN in controlling NaCl preference

Structure-function analyses of *PTEN* variants necessitate an *in vivo* phenotypic assay that reports on *PTEN* function. When wild-type worms are grown in the presence of NaCl and their food source (*Escherichia coli*) they display naïve attractive navigation behavior toward NaCl. This behavior is called NaCl chemotaxis and can be quantified by measuring navigation behavior of a population of animals up a controlled NaCl concentration gradient (Ward, 1973). Previous work has shown that *daf-18* is required for attractive navigation behavior up a concentration gradient of NaCl such that animals with deletion or reduction-of-function mutations in *daf-18* display innate *aversion* to NaCl (Tomioka et al., 2006). Interestingly, Insulin/PI3K signaling normally *actively promotes salt avoidance* under naive conditions and *daf-18* functions in a single chemosensory neuron (ASER) to antagonize the Insulin/PI3K pathway and promote salt attraction (Adachi et al., 2010; Tomioka et al., 2006). Using our machine vision system, the Multi-Worm Tracker, we developed an automated high-throughput NaCl chemotaxis paradigm (Fig 3A,B) and replicated the finding that *daf-18(e1375)* reduction-of-function mutants display strong aversion to NaCl (Swierczek et al. 2011and Fig. 3C,D). We then generated a transgenic line using traditional extrachromomosal array technology that directed pan neuronal expression of wild-type human *PTEN* using the *aex-3* promoter (Kuroyanagi et al., 2010). Pan neuronal expression of human *PTEN* is able to rescue the *daf-18* reduction of function phenotype and restore attractive NaCl chemotaxis to wild-type levels (Fig 3C,D). This work establishes NaCl chemotaxis as an *in vivo* behavioral assay of conserved nervous system *PTEN* functions.

**Figure 3.**
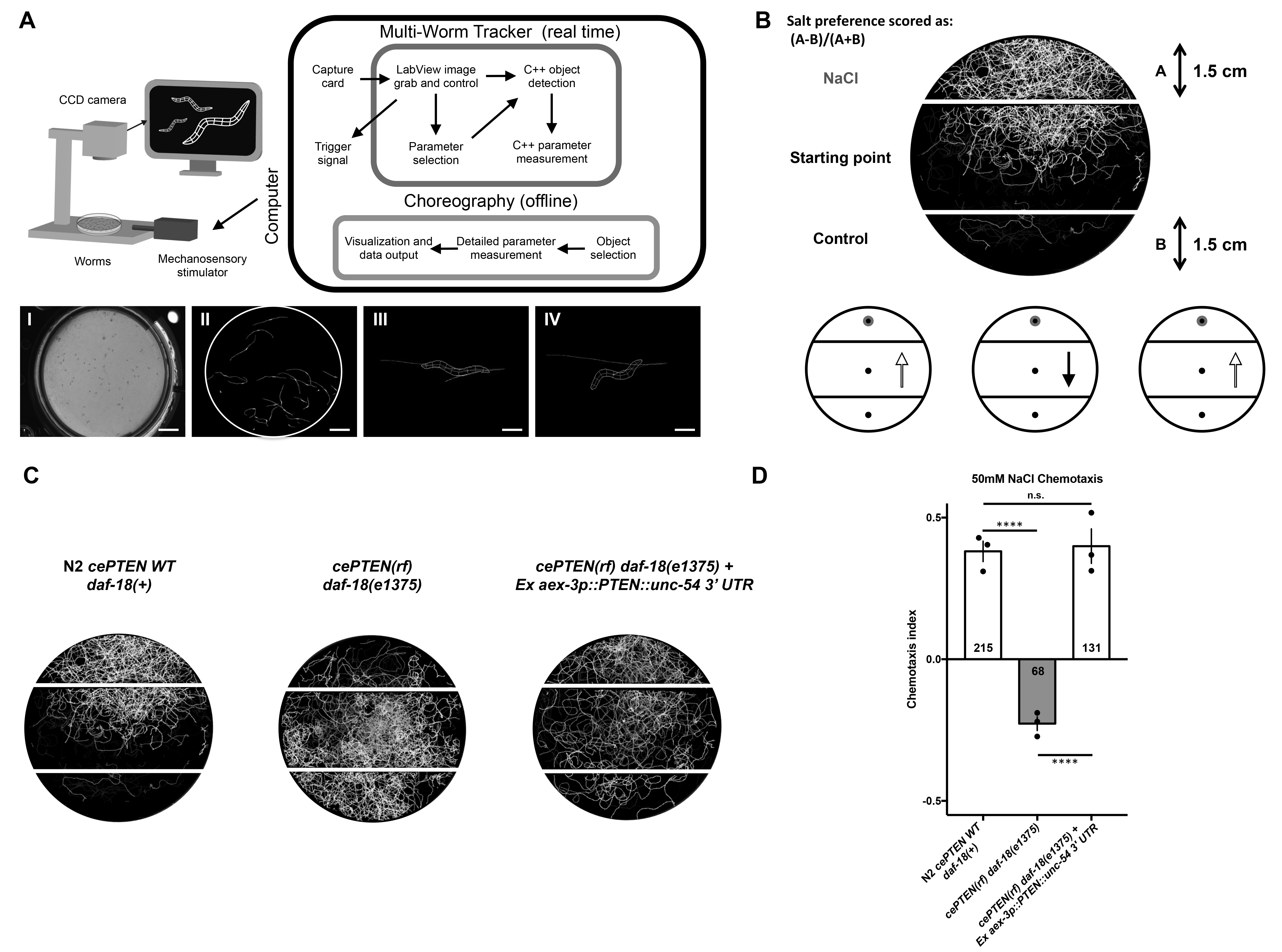
A conserved neuronal role for *PTEN* in NaCl preference revealed by a novel automated chemotaxis paradigm. **A)** (top) All phenotypic analysis is conducted using our machine vision system the Multi-Worm Tracker. The Multi-Worm Tracker delivers stimuli and preforms image acquisition, object selection, and parameter selection in real time while choreography software extracts detailed phenotypic information offline (A bottom panels) I) petri plate of *C. elegans* II) A petri plate of *C. elegans* selected for analysis by the Multi-Worm Tracker III) A Multi-worm tracker digital representation showing the degree of phenotypic detail. An example behaviour scored by the Multi-Worm Tracker: the *C. elegans* response to a mechanosensory tap to the side of the Petri plate is brief backwards locomotion (from III to IV). B) Behavioural track tracing of a plate of worms from a novel Automated Multi-Worm Tracker NaCl chemotaxis paradigm illustrating attractive navigation behaviour of wild-type animals toward a point source of NaCl. **B)** (bottom left to right) circles and arrows and **C)** (left to right) worm tracks represent navigation trajectories of wild-type attraction to a point source of NaCl, a *daf-18(e1375)* reduction-of-function decrease in NaCl chemotaxis, and a transgenic rescue of NaCl preference via pan neuronal overexpression of WT human *PTEN* in *daf-18(e1375)* reduction of function mutants. **D)** Quantitative chemotaxis index scores across genotypes. Pan neuronal expression of human *PTEN* rescues the reversed NaCl preference of *daf-18(e1375)* mutants to wild-type levels. Circles represent plate replicates run on the same day and inset number represent the number of individual animals registered by the tracker and located outside the center starting region (i.e. included in preference score) across the three plate replicates for each genotype. Error bars represent standard deviation using the number of plates as n (n = 3). (****) P < 0.0001, n.s. not significant, one-way ANOVA and Tukey’s post-hoc test.

### Complete deletion of the *daf-18* ORF causes strong NaCl avoidance that is not rescued by direct single-copy replacement with the canonical human *PTEN* CDS

Next we used our strategy to create a *daf-18* deletion allele and single-copy replacement *daf-18* with the 1212 bp canonical human *PTEN* CDS. Complete deletion of the 4723 bp *daf-18* open reading frame resulted in strong aversion to NaCl and chemotaxis down the salt gradient (Fig. 4A). Chemotaxis avoidance of worms harboring the *daf-18* complete deletion allele generated using CRISPR was not significantly different from that of worms carrying the previously characterized large deletion allele *daf-18(ok480)*, confirming effective inactivation of *daf-18* (Fig. 4B). DMS cassette excision and expression of a single copy of human *PTEN* was unable to substitute for *daf-18* and did not rescue attractive NaCl chemotaxis behavior (Fig. 4C). This result was observed in two independent single-copy human *PTEN* knock in lines on several independent experimental replications (Fig. 4B). Transcription of human *PTEN* was confirmed using Reverse transcription PCR in both knock in lines (Fig. 4D). Sanger sequencing confirmed error free insertion of transgenes at base pair resolution before and after cassette excision. There are several potential reasons why expression of human *PTEN* using extrachromasomal arrays rescued *daf-18* mutant phenotypes (Solari et al. 2005 and Fig. 3C,D) while targeted single-copy replacement with *PTEN* did not (Fig. 4C). The two most prominent differences between these two technologies are the expression level of the transgenes and the use of the endogenous 3’ UTR in the CRISPR knock in versus the *unc-54* myosin 3’ UTR used in most *C. elegans* transgenes to ensure proper processing of transcripts in all tissues, including all constructs previously shown to rescue *daf-18* phenotypes with human *PTEN* (Merritt and Seydoux, 2010; Solari et al., 2005, and Fig. 3C,D).

**Figure 4.**
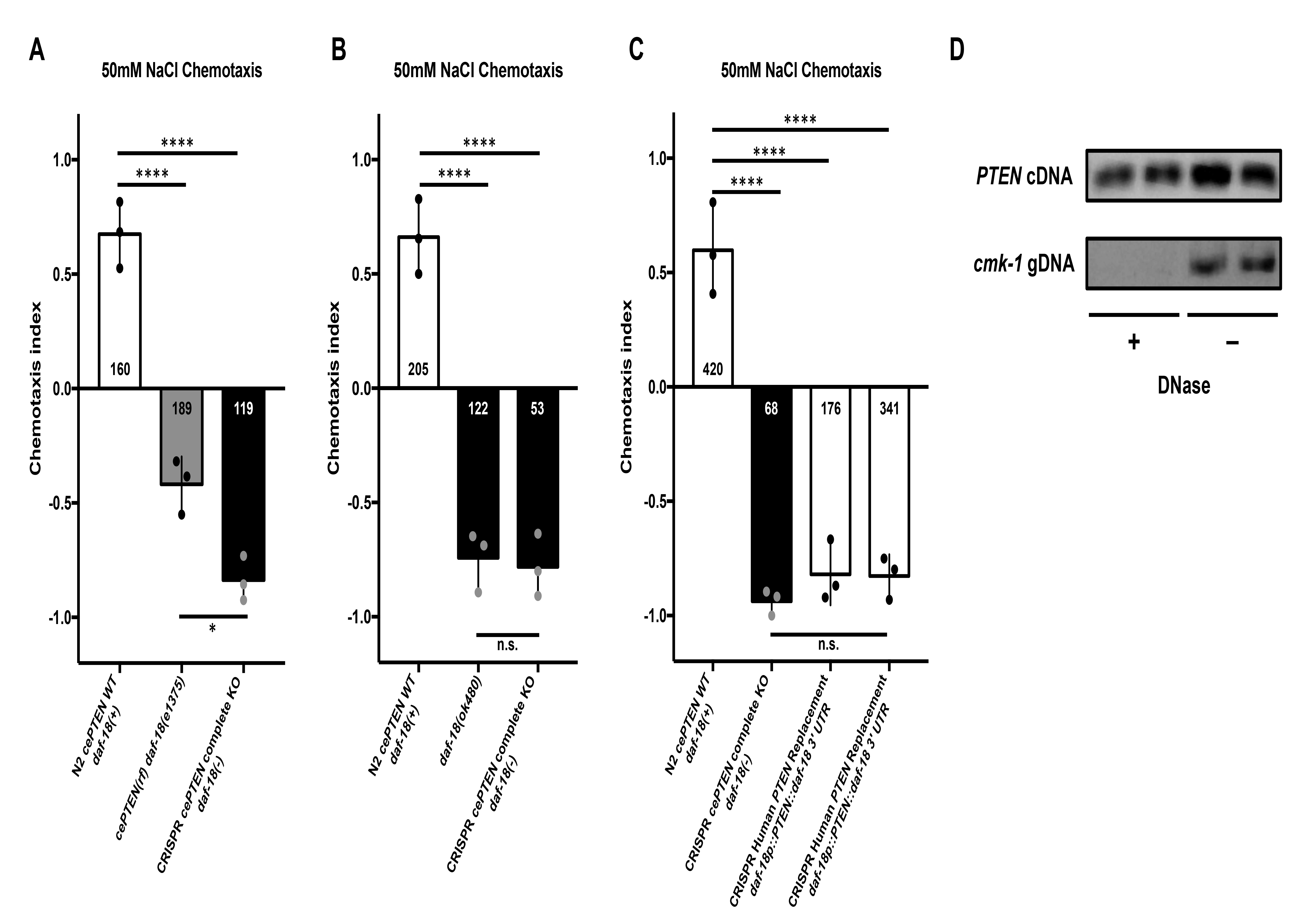
Complete *daf-18* ORF deletion causes strong NaCl avoidance that is not rescued by direct replacement with human *PTEN*. **A)** NaCl chemotaxis preference scores of wild-type, *daf-18(e1375)* reduction of function, and CRISPR *daf-18* complete deletion mutants. *daf-18* complete deletion mutants show significantly stronger NaCl avoidance than *daf-18(e1375)* reduction of function mutants. **B)** CRSPR *daf-18* complete deletion mutants are not significantly different from animals harboring the putative null *daf-18(ok480)* deletion allele. **C)** Expression of a single copy *PTEN* transgene from the native *daf-18* locus is insufficient to significantly rescue NaCl chemotaxis towards wild-type levels. **D)** RT-PCR confirming expression of full length PTEN mRNA in the two independent knock-in lines used for behavioural analysis. Previously validated primers that target *cmk-1* intronic regions of genomic DNA do not produce products following DNase treatment, confirming purity of the cDNA. Circles represent plate replicates run on the same day and inset number represent the number of individual animals registered by the tracker and located outside the center starting region (i.e. included in preference score) across the three plate replicates for each genotype. Error bars represent standard deviation using the number of plates as n (n = 3). (****) P < 0.0001, (*) P < 0.05 n.s. not significant, one-way ANOVA and Tukey’s post-hoc test.

### A streamlined human gene replacement strategy functionally replaces *daf-18* with human *PTEN*

In order to address these limitations and increase the speed of transgenesis we created an alternate repair template strategy that includes a transcriptional termination sequence in the upstream homology arm so that expression of the human gene begins immediately upon integration (Fig. 5A). We included the validated *unc-54 3’ UTR* sequence in the upstream homology arm fused to the 3’ end of human *PTEN* (Fig. 5A). Expression of wild-type human *PTEN* using this genome editing strategy rescued NaCl chemotaxis, indicating production of functional PTEN immediately upon genomic integration prior to cassette excision (Fig. 5B). This alternate approach also offers the added benefits of increased throughput (as a second injection step to excise the cassette was not required for human gene expression as it is in the first strategy) and the option for retained visual transgenic markers (either within the selection cassette or by adding a 2A sequence to drive reporter expression from the same promoter), which simplifies the generation and phenotypic analysis of heterozygotes and double mutants (Fig. 5A, Ahier and Jarriault, 2014; Calarco and Norris, 2018; Norris et al., 2017). Given the demonstrated versatility of CRISPR genome editing in *C. elegans*, these results suggest our strategy should be broadly applicable for *in vivo* analysis of diverse human disease-associated genes.

**Figure 5.**
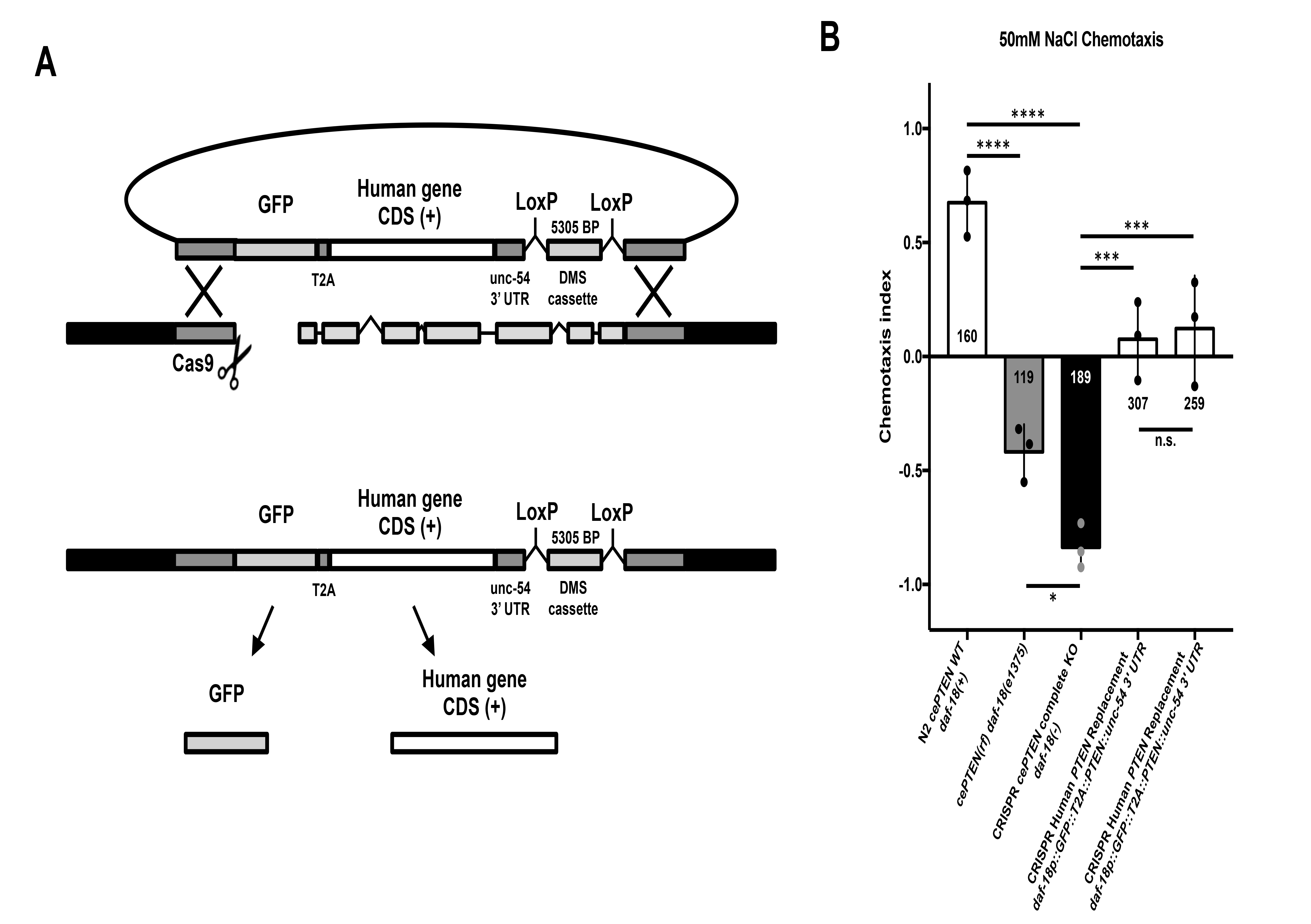
A streamlined human gene replacement strategy functionally replaces *daf-18* with human *PTEN*. **A)** Streamlined CRISPR gene replacement strategy. Inclusion of the validated *unc-54* 3’ UTR in the upstream homology arm increases the speed of transgenesis by removing the need for cassette excision to induce transgene expression. This alternate approach also offers the option for retained visual transgenic markers, which simplifies the generation and phenotypic analysis of heterozygotes and double mutants. The inclusion of a GFP::T2A cassette is an optional addition to allow for confirmation of transgene expression without altering human gene function (Ahier and Jarriault, 2014). **B)** Expression of wild-type human *PTEN* using this genome editing strategy significantly rescued NaCl chemotaxis toward wild-type levels. Circles represent plate replicates run on the same day and inset number represent the number of individual animals registered by the tracker and located outside the center starting region (i.e. included in preference score) across the three plate replicates for each genotype. Error bars represent standard deviation using the number of plates as n (n = 3). (****) P < 0.0001, (***) P < 0.001, (*) P < 0.05, n.s. not significant, one-way ANOVA and Tukey’s post-hoc test.

### *daf-18* deletion causes mechanosensory hyporesponsivity that is rescued by targeted replacement with human *PTEN*

A long-standing goal has been to understand how human disease-associated variants alter normal gene function to produce sensory and learning abnormalities characteristic of diverse neurogenetic disorders. The massive number variants of uncertain significance recently implicated in the etiology neurogenetic disorders necessitates a dramatic increase in throughput of both transgenic construction and behavioral phenoptying if this goal is to be achieved (Ben-Shalom et al., 2017; Geschwind and Flint, 2015; Lim et al., 2017; Sanders et al., 2015; Starita et al., 2017; Wangler et al., 2017). By combing streamlined human gene integration with rapid machine vision phenotypic analysis of *C. elegans* our strategy greatly simplifies the identification of novel conserved gene functions in complex sensory and learning phenotypes.

We explored whether *daf-18* mutants displayed behavioral deficits in mechanosensory responding and/or habituation, a conserved form of non-associative learning that is altered in several neurodevelopmental and neuropsychiatric disorders (McDiarmid et al., 2018, 2017; Rankin et al., 2009; Stessman et al., 2017; Timbers et al., 2017; van der Voet et al., 2014). When a non-localized mechanosensory stimulus is delivered to the side of the petri plate *C. elegans* are cultured on they respond by eliciting a brief reversal before resuming forward locomotion. Wild-type *C. elegans* habituate to repeated stimuli by learning to decrease the probability of eliciting a reversal (Rankin et al., 1990). To determine if *daf-18* is important for mechanoresponding and/or non-associative learning we examined habituation of the *daf-18(e1375)* and *daf-18* complete deletion mutants. Compared to wild-type animals both *daf-18(e1375)* reduction of function and *daf-18* complete deletion mutants exhibited significantly reduced probability of eliciting a reversal response throughout the habituation training session, indicating mechanosensory hyporesponsivity (Fig. 6A,B). Despite this hyporesponsivity, the plasticity of responses, or the pattern of the gradual decrement in the probability of emitting of a reversal response throughout the training session was not significantly altered in *daf-18* mutants (Fig. 6A,B,E). Importantly, targeted single-copy replacement of *daf-18* with human *PTEN* was sufficient to rescue the mechanosensory hyporesponsivity phenotype across the training session towards wild-type levels (Fig. 6D). These results identify a novel conserved role for *PTEN* in mechanosensory responding, a fundamental biological process disrupted in diverse genetic disorders (Badr et al., 1987; McDiarmid et al., 2017; Orefice et al., 2016). More broadly, they illustrate how the library of transgenic animals generated with our strategy can be used to rapidly characterize the role of diverse human genes in complex sensory and learning behaviors. These novel phenotypes can then be used to investigate the functional consequences of disease-associated variants in intact and freely behaving animals and to screen for therapeutics that reverse the effects of a particular patients missense variant.

**Figure 6.**
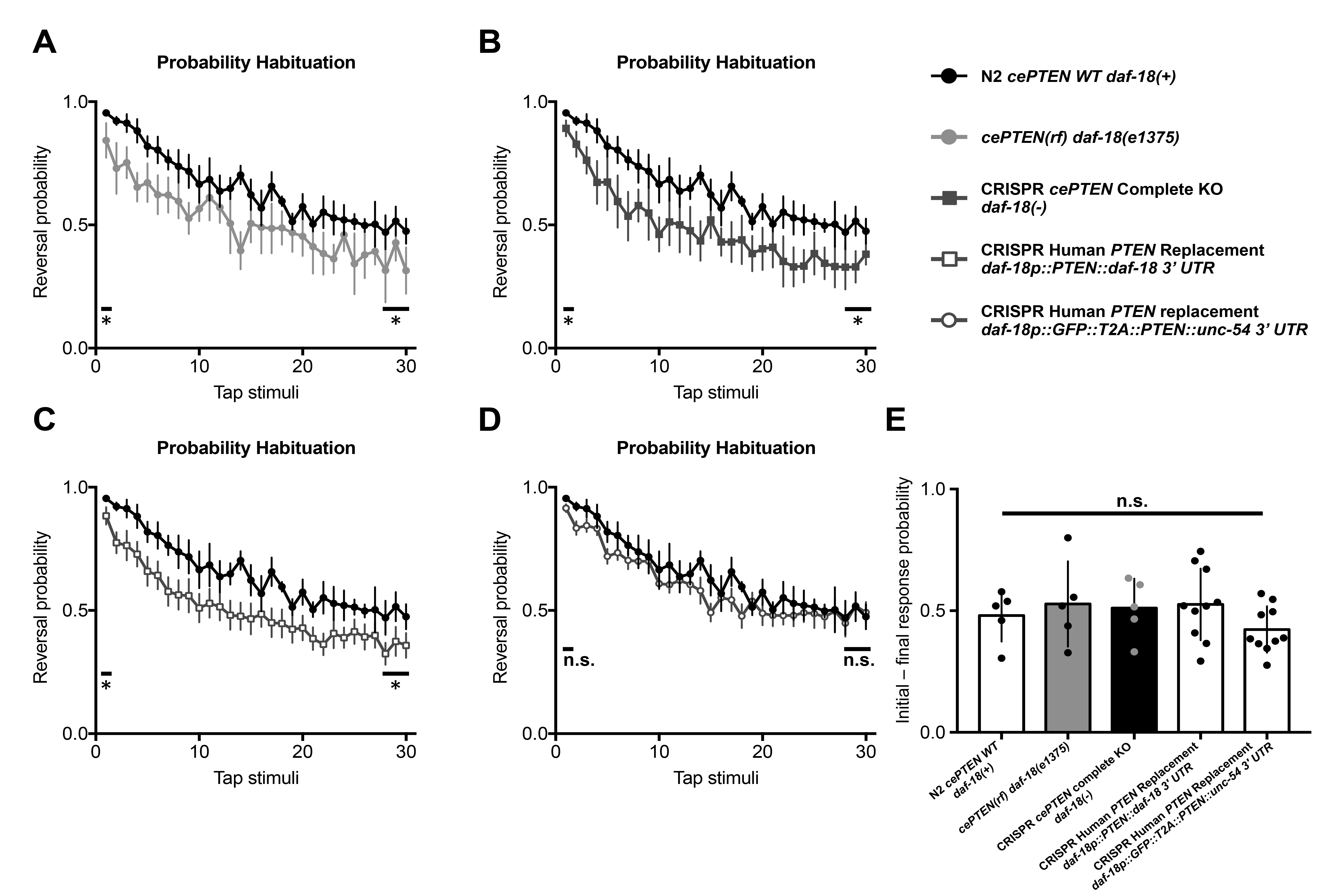
*daf-18* deletion causes mechanosensory hyporesponsivity that is rescued by targeted replacement with human *PTEN*. **A-D)** Average probability of eliciting a reversal response to 30 consecutive tap stimuli delivered at a 10s ISI. A) *daf-18(e1375)* reduction of function and **B)** *daf-18* complete deletion mutants exhibit significantly reduced probability of eliciting a reversal response throughout the habituation training session, indicating mechanosensory hyporesponsivity. **C)** Replacement of *daf-18* with human *PTEN* using the original strategy does not rescue mechanosensory hyporesponsivity. **D)** Expression of human *PTEN* using the streamlined replacement strategy rescues mechanosensory responding to wild-type levels. Error bars represent standard error of the mean. **E)** Habituation, or the ability to learn to decrease the probability of eliciting of a reversal response throughout the training session was not significantly altered in *daf-18* mutants. Circles represent plate replicates run on the same day. Error bars represent standard deviation of the mean using the number of plates as n (n = 5 or 10). (*) P < 0.05 n.s. not significant, one-way ANOVA and Tukey’s post-hoc test.

### Human gene replacement and *in vivo* phenotypic assessment accurately identifies functional consequences of the pathogenic PTEN-G129E variant

To demonstrate the feasibility of our human gene replacement strategy for assessing variants of uncertain significance we set out to determine whether our *in vivo* functional assays could discern known pathogenic variants. Early studies characterizing the role of PTEN as a tumor suppressor suggested impaired protein phosphatase activity was key to the etiology of PTEN disorders (Tamura et al., 1998). However, this notion was challenged by the identification of cancer patients harboring a missense mutation that changes a glycine residue in the catalytic signature motif to a glutamate, which was predicted to abolish the lipid phosphatase activity of PTEN while leaving the protein phosphatase activity intact (Liaw et al., 1997; Myers et al., 1997). Subsequent biochemical analyses supported the now widely accepted view that the lipid phosphatase activity of PTEN is critical to its tumor suppressor activity (Myers et al., 1998).

We used site-directed mutagenesis to incorporate the PTEN-G129E (*PTEN*, c386G>A) missense variant into our repair template and used our human gene replacement strategy to replace *daf-18* with a single copy of human PTEN-G129E (Fig. 7A). Animals harboring the PTEN-G129E variant displayed strong NaCl avoidance equivalent to animals carrying the complete *daf-18* deletion allele, indicating loss-of-function (Fig. 7B). Similarly, PTEN-G129E mutants also displayed mechanosensory hyporesponsivity that was not significantly different from *daf-18* deletion carriers (Fig. 7C). These *in vivo* phenotypic results accurately classify the pathogenic PTEN-G129E as a strong loss-of-function variant. In addition, by taking advantage of a pathogenic variant with well-characterized biochemical effects these results identify a necessary role for PTEN lipid phosphatase activity in both chemotaxis and mechanosensory responding, providing novel insight into the molecular mechanisms underlying these forms of sensory processing. Taken together, these results demonstrate the potential of human gene replacement and phenomic characterization to rapidly identify the functional consequences variants of uncertain significance.

**Figure 7.**
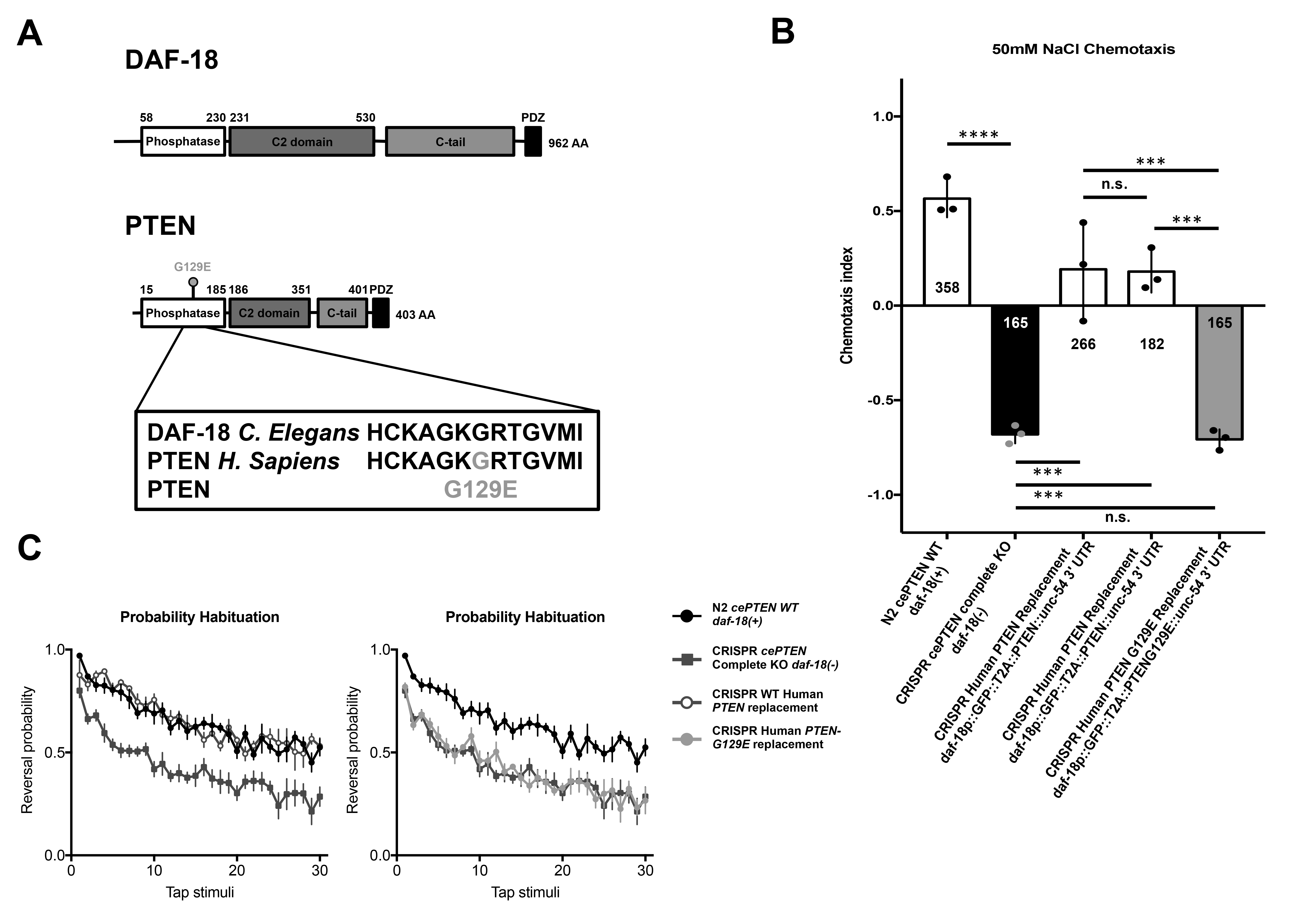
Human gene replacement and *in vivo* phenotypic assessment accurately identifies functional consequences of the pathogenic PTEN-G129E variant. **A)** Location (top) and conserved amino acid sequence change (bottom) of the pathogenic lipid phosphatase inactive PTEN-G129E variant within the PTEN phosphatase domain. **B)** Animals harboring the PTEN G129E variant displayed strong NaCl avoidance equivalent to animals carrying the complete *daf-18* deletion allele, indicating loss-of-function (Fig. 7B). Circles represent plate replicates run on the same day and inset number represent the number of individual animals registered by the tracker and located outside the center starting region (i.e. included in preference score) across the three plate replicates for each genotype. Error bars represent standard deviation using the number of plates as n (n = 3). **C)** Similarly, PTEN-G129E mutants also displayed mechanosensory hyporesponsivity that was not significantly different from *daf-18* deletion mutants. Error bars represent standard error of the mean. (****) P < 0.0001, (***) P < 0.001, (*) P < 0.05, n.s. not significant, one-way ANOVA and Tukey’s post-hoc test.

## Discussion

We have developed and validated a broadly applicable strategy for targeted human gene replacement and phenomic characterization in *C. elegans* that will facilitate assessment of the functional conservation of human genes and structure-function analysis of variants of uncertain significance with unprecedented precision. We established an automated NaCl chemotaxis paradigm and demonstrate that pan neuronal overexpression or direct replacement of *daf-18* with its human ortholog *PTEN* using CRISPR is sufficient to rescue reversed NaCl chemotaxis preference induced by complete *daf-18* deletion. We further identified a novel mechanosensory hyporesponsive phenotype for *daf-18* mutants that could also be rescued by targeted replacement with human *PTEN*. *In vivo* characterization of mutants harboring a single copy of the known lipid phosphatase inactive G129E variant accurately classified this variant as pathogenic and revealed a critical role for PTEN lipid phosphatase activity in NaCl chemotaxis and mechanosensory responding. As a resource article, we provide novel high-throughput *in vivo* functional assays for *PTEN*, as well as validated strains, reagents, sgRNA and repair template constructs to catalyze further analysis of this critical human disease-associated gene. More broadly, we provide a conceptual framework that illustrates how genome engineering and automated machine vision phenotyping can be combined to efficiently generate and characterize a library of knockout and humanized transgenic strains that will allow for straightforward and precise analysis of human genes and disease-associated variants *in vivo* (Fig. 8).

**Figure 8.**
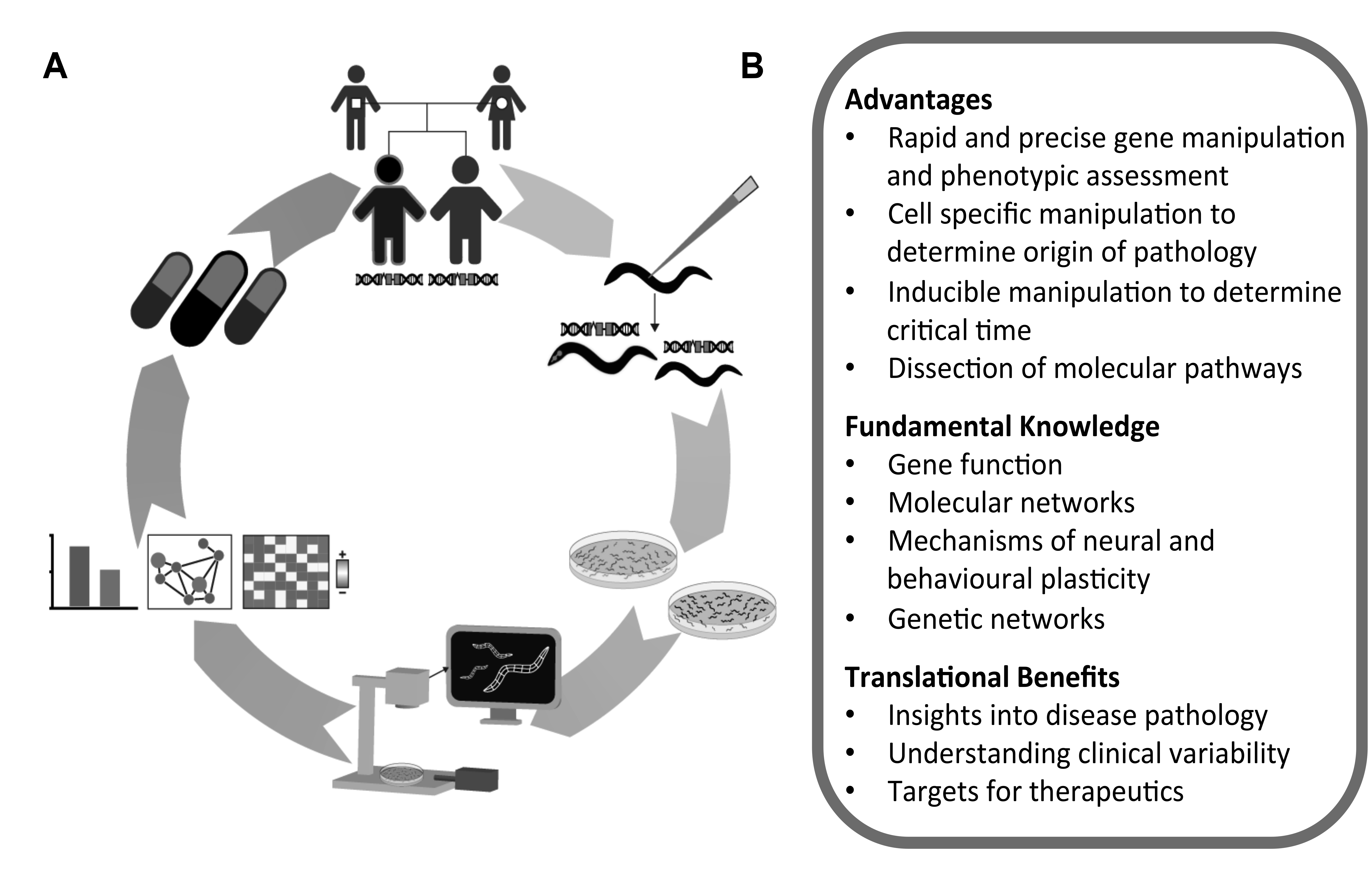
A conceptual framework for *in vivo* functional analysis of human genetic variation using *C. elegans*. **A)** (top working clockwise) A human gene and/or variant of uncertain significance is implicated in disease etiology through clinical sequencing. Targeted CRISPR human gene replacement or analogous methods are used to generate a library of knock-out, human wild-type and variant transgenic strains. Large isogenic synchronous colonies of these transgenic worms are grown and their morphology, baseline locomotion, and sensory phenotypes are rapidly characterized using machine vision to establish novel functional assays and interpret variant effects. In vivo functional data can be used to probe epistatic network disruptions and cluster variants based on multi-parametric phenotypic profiles. The integrated humanized transgenic lines and functional assays greatly facilitate downstream applications including precision medicine drug screens designed to identify compounds that reverse the effects of a particular patients missense variant. **B)** Advantages of targeted human gene replacement using *C. elegans.*

### Comparing CRISPR targeted human gene replacement with orthology-based variant assessment methods

To date, the most widely used genome editing-based human disease variant assessment strategy in *C. elegans* uses sequence alignments to identify and engineer the corresponding amino acid change into the orthologous *C. elegans* gene (Bend et al., 2016; Bulger et al., 2017; Canning et al., 2018; Pierce et al., 2018; Prior et al., 2017; Sorkaç et al., 2016; Troulinaki et al., 2018). A major advantage of this approach is that by using the *C. elegans* gene, intronic regulation, protein-protein interactions, subcellular localization, and biochemical activity of the protein of interest are, by design, perfectly modeled by *C. elegans*. Even when expressed at physiologically relevant levels directly from the orthologs native loci, and with evidence of phenotypic rescue, it is not guaranteed a transgenic human protein will recapitulate all functions and interactions of the orthologous *C. elegans* protein. This presents an important consideration when attempting to replace *C. elegans* proteins that must interact in extremely precise heteromeric complexes to perform their molecular functions (e.g. certain ion channel subunits) (Bend et al., 2016; Prior et al., 2017). The most obvious limitation of this approach is that it can only be used to study orthologous amino acids. Amino acids that have been conserved throughout evolution are, by definition, the least tolerant to mutation and are thus far more likely to be detrimental to protein function when mutated (Starita et al., 2017; Weile et al., 2017). Indeed, many variant effect prediction algorithms rely on sequence conservation as the main predictor that a variant will be deleterious (Richards et al., 2015; Starita et al., 2017). It is also important to note that a large portion of amino acids will not be conserved to humans (e.g. >50% of amino acids are not conserved from DAF-18 to PTEN, Fig. 2C) and alignment algorithms that identify orthologous amino acids are imperfect. Many current implementations of orthologous amino acid engineering also require constant generation of completely new sgRNAs and repair templates for subsequent edits. Human gene replacement, in contrast, allows any coding variant of uncertain significance to be studied *in vivo* using the same validated sgRNA and homology arms.

One potential limitation of human gene replacement is that it relies on a *C. elegans* ortholog to replace. Depending on the orthology prediction program used estimates suggest there are *C. elegans* orthologs for 60-80% of human disease-associated genes meaning their will not be an ortholog available for a minority of human genes (Kaletta and Hengartner, 2006; Lai et al., 2000b; Shaye and Greenwald, 2011). This is even less likely to be a problem for disease variant modelling, as disease-associated genes are more likely to be highly conserved (Aerts et al., 2006; López-Bigas and Ouzounis, 2004). A recent study also showed that ~10% of human disease-associated genes are able to functionally substitute for their yeast paralogs (in addition to orthologs), further increasing the number of human genes that can be studied by replacement (Yang et al., 2017). Still, in the event there is no suitable *C. elegans* ortholog or paralog available for a human gene of interest the human gene can simply be integrated at a putatively neutral genomic location using CRISPR or transposon mobilization and expressed from a heterologous promoter (Frøkjær-Jensen et al., 2014, 2012, 2008). This approach can be used to screen for phenotypes induced by expression of any human gene and to determine whether a variant exacerbates or eliminates these effects (Baruah et al., 2017).

With the rapidly expanding set of precise genome editing techniques available to *C. elegans,* researchers interested in variants of uncertain significance now have the freedom to choose the approach that best suits their particular needs and interests. The approaches described here provide a diverse collection of methods that can be sequentially tested in a pragmatic hierarchy of precision, beginning with direct replacement and working down until phenotypic rescue is achieved. Regardless, it will always be ideal to have corroborating evidence of variant effect from multiple techniques, indeed multiple model systems, to best inform clinical decisions.

### Combining human gene replacement and automated phenomic characterization to discover conserved gene functions and establish variant functional assays

A necessary step in the establishment of human gene functional assays is the identification of phenotypes that are rescued by or induced upon expression of the reference (wild-type) human gene, as we have done here for human *PTEN* and NaCl chemotaxis. Indeed, the establishment of functional assays remains a major bottleneck for variant assessment across species (Starita et al., 2017; Weile et al., 2017). While traditional extrachromosomal array transgenes offer a quick way to establish such assays several aspects of these transgenes can severely impede this process. These include but are not limited to: 1) variable overexpression of transgenes which can lead to silencing of transgenes in certain tissues, complicating phenotypic analysis (e.g. multi-copy transgenes are expressed in the soma but silenced in the germline while low-and single-copy transgenes are expressed in both) (Kelly et al., 1997; Merritt and Seydoux, 2010), and 2) variably mosaic expression which can make it extremely difficult to assess rescue of partially penetrant and subtle complex phenotypes, as an animal must simultaneously carry the extrachromosomal array and be one of the members of the population that displays the partially penetrant phenotype that can often be difficult to score.

The human gene replacement approach described here allows for generation of ortholog deletion alleles directly in any wild-type or mutant strain amenable to transgenesis using the same reagents designed for replacement, thereby reducing the confounding effects of background mutations on phenotype discovery. The use of excisable selectable markers that do not severely alter morphology, baseline locomotion, and several evoked sensory behaviors further simplifies phenotypic analysis and removes the need for any specialized genetic backgrounds (Norris et al. 2015 and Figs 4, 5, 6, S1). Using this approach in combination with machine vision we provide two *in vivo* functional assays for human *PTEN*, NaCl chemotaxis and mechanosensory responsivity. In particular, NaCl chemotaxis possesses several characteristics that make it an ideal functional assay: 1) a large functional range between deletion and human gene rescue to discern potentially subtle functional differences among missense variants; 2) scalability as many plates can be run simultaneously; 3) straightforward analysis using an automatically calculated preference index (alternatively many labs score chemotaxis manually by simple blinded counting). Importantly, these reagents and functional assays can now be used in precision medicine drug screens aimed at identifying compounds that counteract the affect of a particular patient’s missense variant. Further broad-scale phenomic characterization of targeted knockouts and mutant libraries combined with new databases that curate ortholog functional annotation across model organisms should expedite the process of *in vivo* functional assay development (Lee et al., 2018; McDiarmid et al., 2018; Thompson et al., 2013; Wang et al., 2017; Yemini et al., 2013).

### Further applications of targeted human gene replacement

One of the goals of this resources article is to illustrate how Cas9-triggered homologous recombination can be adapted to directly replace *C. elegans* genes with human genes. An exciting adaptation of this approach will be to combine targeted human gene insertions with bi-partite systems for precise spatial-temporal control of human transgene expression, as has recently been done to overexpress human genes as UAS-cDNA constructs in *Drosophila* (Luo et al. 2017; Lee et al. 2018; Wangler et al. 2017). The recent and long-awaited development of the cGAL4-UAS system should allow a similar approach to be developed for *C. elegans*, currently the only organism where the complete cell lineage and neuronal wiring diagram is known (Sulston et al., 1983; Sulston and Horvitz, 1977; Wang et al., 2016; White et al., 1986).

While one of the clearest uses for targeted replacement is precise structure-function analysis of variants of uncertain significance there are several exciting applications beyond modeling disease-associated alleles. Targeted human gene replacement is also particularly well suited for investigation of the evolutionary principles that determine the replaceability of genes. By allowing for human gene expression immediately upon genomic integration this approach should be also adaptable to essential genes. A homozygous integrant will only be obtained if the human gene can substitute for the orthologous essential gene, creating a complementation test out of the transgenesis process. This will allow for systematic and precise interrogation of the sequence characteristics and functional properties required for successful human gene replacement (Kachroo et al., 2015). A library of humanized worms would also open the door to the rich resources of tools available to visualize and manipulate human genes and proteins that are often unavailable for *C. elegans* researchers (e.g. high-quality antibodies, biochemically characterized or known pathogenic control variants, and experimentally determined crystal structures, Figs 2, 7; Berman et al. 2000). Given the throughput that has been achieved for reporter gene analysis (several thousand genes) and genome editing in *C. elegans* it should be possible to generate a humanized *C. elegans* library of similar size to those recently created in yeast (Dickinson and Goldstein, 2016; Dupuy et al., 2007; Hamza et al., 2015; Hunt-Newbury et al., 2007; Kachroo et al., 2015; Norris et al., 2017; Sun et al., 2016; Yang et al., 2017). Integrated transgenes would offer the possibility of humanizing entire cellular processes for detailed *in vivo* analysis in a relatively complex yet tractable metazoan with increasingly powerful tools for spatial-temporal control of transgene expression and protein degradation (Armenti et al., 2014; Wang et al., 2016; S. Wang et al., 2017; Zhang et al., 2015).

Deep mutational scanning and related technologies have recently made it feasible to characterize the functional effects of virtually every possible amino acid change of a protein on a particular cellular phenotype (Fowler et al., 2010; Fowler and Fields, 2014). Several such exhaustive sequence-function maps have been recently been generated in yeast and human cell culture systems (Findlay et al., 2018, 2014; Majithia et al., 2016; Matreyek et al., 2018; Mighell et al., 2018; Weile et al., 2017). These tools offer amazing resources that serve as ‘lookup tables’ of functional missense variation in human genes, to enable experimentally confirmed variant interpretation immediately upon first clinical presentation (Starita et al., 2017; Weile et al., 2017). An ambitious and exciting goal for the *C. elegans* community will be to further streamline genome engineering and high-throughput phenotyping to achieve the first comprehensive sequence-function map in a metazoan.

## Materials and Methods

### Strains and culture

Worms were cultured on Nematode Growth Medium (NGM) seeded with *Escherichia coli* (OP50) as described previously (Brenner, 1974). N2 Bristol, and CB1375 *daf-18(e1375)* strains were obtained from the *Caenorhabditis* Genetics Center (University of Minnesota, USA). *daf-18(e1375)* harbors a 30–base pair insertion in the fourth exon and is predicted to insert six amino acids before introducing an early stop codon that truncates the C-terminal half of the protein while leaving the phosphatase domain intact (Ogg and Ruvkun, 1998).

The following strains were created for this work:

VG674 *daf-18(e1375); yvEx674[paex-3::PTEN::unc-54; pmyo-2::mCherry::unc-54 UTR]*

VG810-813 *daf-18(e1375); yvEx810-813[paex-3::PTEN::unc-54 UTR; pmyo-2::mCherry::unc-54 UTR]*

VG712 *daf-18(yv3[daf-18p::PTEN + LoxP pmyo-2::GFP::unc-54 UTR prps-27::NeoR::unc-54 UTR LoxP + daf-18 UTR])*

VG713 *daf-18(yv4[daf-18p::PTEN + LoxP pmyo-2::GFP::unc-54 UTR prps-27::NeoR::unc-54 UTR LoxP +daf-18 UTR])*

VG714 *daf-18(yv5[daf-18p::PTEN + LoxP + daf-18 UTR])*

VG715 *daf-18(yv5[daf-18p::PTEN + LoxP + daf-18 UTR])*

VG817 *daf-18(yv7[daf-18p::GFP::T2A::PTEN::unc-54 UTR + LoxP pmyo-2::GFP::unc-54 UTR prps-27::NeoR::unc-54 UTR LoxP + daf-18 UTR])* VG818 *daf-18(yv8[daf-18p::GFP::T2A::PTEN::unc-54 UTR + LoxP pmyo-2::GFP::unc-54 UTR prps-27::NeoR::unc-54 UTR LoxP + daf-18 UTR])*

VG867 *daf-18(yv14[daf-18p::GFP::T2A::PTEN*-G129E*::unc-54 UTR + LoxP pmyo-2::GFP::unc-54 UTR prps-27::NeoR::unc-54 UTR LoxP + daf-18 UTR])*

### Strain and plasmid generation

The reference *PTEN* CDS (UniProt consensus, identifier: P60484-1) was obtained from a pCMV-PTEN plasmid (Addgene plasmid #28298) and cloned into a TOPO gateway entry clone (Invitrogen) according to manufacturers instructions. The *PTEN* entry clone was recombined with an pDEST-*aex-3p* destination vector (obtained from Dr. Hidehito Kuroyanagi; Kuroyanagi et al., 2010) to generate the *aex-3p::PTEN::unc-54 UTR* rescue construct using gateway cloning (Invitrogen), according to manufacturers instructions.

The Moerman lab guide selection tool (http://genome.sfu.ca/crispr/) was used to identify the *daf-18* targeting sgRNA. The *daf-18* sgRNA sequence: GGAGGAGGAGTAACCATTGG was cloned into the *pU6::klp-12* sgRNA vector (obtained from Calarco lab) using site-directed mutagenesis and used for all editing experiments. The *daf-18p::PTEN* CDS and *daf-18p::GFP::T2A::PTEN::unc-54 UTR* upstream homology arms were synthesized by IDT and cloned into the loxP_myo2_neoR repair construct (obtained from Calarco lab) using Gibson Assembly.

*C. elegans* wild-type N2 strain was used for all CRISPR editing experiments. Genome edits were created as previously described (Norris et al., 2015). In brief, plasmids encoding sgRNA, Cas9 co-transformation markers pCFJ90 and pCFJ104 (Jorgensen lab, Addgene) and the selection cassette flanked by homology arms (~500 bp) containing *PTEN* were injected into wild-type worms. Animals containing the desired insertions were identified by G418 resistance, loss of extrachromosomal array markers, and uniform dim fluorescence of the inserted GFP.

### Genotype confirmation

Correct replacement of the *daf-18* ORF with human *PTEN* was confirmed by amplifying the two regions spanning the upstream and downstream insertion borders using PCR followed by Sanger sequencing (primer binding locations depicted in Fig. 1). The genotyping strategy is essentially as described for deletion allele generation via DMS cassette insertion in (Norris et al. 2015).

The forward and reverse primers used to amplify the upstream insertion region were TGCCGTTTGAATTAGCGTGC (located within the *daf-18* genomic promoter region) and CCCTCAATGTCTCTACTTGT (located within the *myo-2* promoter of the selection cassette) respectively.

The forward and reverse primers used to amplify the downstream insertion region were TTCCTCGTGCTTTACGGTATCG (located within the Neomycin resistance gene) and CTCAACACGTTCGGAGGGTAAA (located downstream of the *daf-18* genomic coding region) respectively.

Following cassette excision via injection of cre-recombinase the *daf-18* promoter (TGCCGTTTGAATTAGCGTGC) and *daf-18* downstream (CTCAACACGTTCGGAGGGTAAA) primers were used to amplify human *PTEN* and confirm error free insertion at the *daf-18* locus via Sanger sequencing (Fig. 1).

### RNA extraction, library preparation, and cDNA amplification

Total RNA was isolated from mixed stage VG714 and VG715 *PTEN* knock in animals using a GeneJET RNA Purification Kit (ThermoFisher) according to manufacturers instructions. Total RNA was treated with DNase (New England Biolabs) and purified with an RNeasy MinElute spin column (Qiagen) according to manufacturers instructions. cDNA libraries were prepared from crude and purified total RNA using Superscript III (Invitrogen). All genes were amplified from cDNA libraries with Platinum Taq DNA Polymerase (ThermoFisher) and gene-specific primer sets.

The forward and reverse primers used to amplify the *PTEN* CDS were ATGACAGCCATCATCAAAGA and TCAGACTTTTGTAATTTGTG respectively.

The forward and reverse *cmk-1* intronic control primers (Ardiel et al., 2018) were AGGGTAGGCTAGAGTCTGGGATAGAT and ACGACTCCGTTGTCGTGCATAAAC respectively.

### Protein structure modeling and visualization

The PTEN 1D5R reference structure (Berman et al., 2000; Lee et al., 1999) was visualized using PyMOL software (DeLano, 2002). Structural models for full-length human PTEN, DAF-18 53-506, and full-length DAF-18 were predicted using Phyre2 (Kelley et al., 2015) and visualized using PyMOL.

### NaCl chemotaxis behavioral assays

The chemotaxis behavioral assay was conducted on a 6 cm assay plate (2% agar), where a salt gradient was formed overnight by inserting a 2% agar plug containing 50mM of NaCl (approximately 5 mm in diameter) 1 cm from the edge of the plate. A control 2% agar plug without NaCl was inserted 1 cm from the opposite edge of the plate. Strains were grown on NGM plates seeded with *E. coli* (OP50) for 3 or 4 days. Worms on the plates were collected and washed three times using M9 buffer before being pipetted onto an unseeded NGM plate to remove excess buffer and select animals carrying transformation markers. Adult worms were transferred and placed at the centre of the assay plates and were tracked for 40 minutes on the Multi-Worm Tracker (Swierczek et al., 2011). After the tracking period, the chemotaxis index was calculated as (A – B)/(A + B), where A was the number of animals that were located in a 1.5 cm-wide region on the side of the assay plate containing the 2% agar plug with 50mM NaCl and B was the number of animals that were located in a 1.5 cm-wide region on the side of the assay plate containing the 2% agar plug without NaCl (Fig. 3B). Animals not located in either region (ie. the middle section of the assay plate) were not counted towards the chemotaxis index. One hundred to two hundred animals were used per plate, and two or three plate replicates were used for each line in each experiment. Any statistical comparisons were carried out on plates assayed concurrently (i.e. on the same day).

### Mechanosensory habituation behavioral assays

Worms were synchronized for behavioral testing on Petri plates containing Nematode Growth Media (NGM) seeded with 50 μl of OP50 liquid culture 12-24 hours before use. Five gravid adults were picked to plates and allowed to lay eggs for 3-4 hours before removal. The animals were maintained in a 20°C incubator for 72 hours. Plates of worms were placed into the tapping apparatus and after a 100s acclimatization period, 30 taps were administered at a 10s ISI. Comparisons of “final response” comprised the average of the final three stimuli. Any statistical comparisons were carried out on plates assayed concurrently (i.e. on the same day).

### Multi-worm tracker behavioral analysis and statistics

Multi-Worm Tracker software (version 1.2.0.2) was used for stimulus delivery and image acquisition (Swierczek et al., 2011). Behavioral quantification with Choreography software (version 1.3.0_r103552) used “--shadowless”, “--minimum-move-body 2”, and “--minimum-time 20” filters to restrict the analysis to animals that moved at least 2 body lengths and were tracked for at least 20 s. The MeasureReversal plugin was used to identify reversals occurring within 1 s (d*t* = 1) of the mechanosensory stimulus onset. Custom MatLab and R scripts organized and summarized Choreography output files. Final figures were generated using GraphPad Prism version 7.00 for Mac OS X. Each experiment was independently replicated at least twice. No blinding was necessary because the Multi-Worm Tracker scores behavior objectively. Morphology metrics, baseline locomotion metrics, initial and final reversal responses, habituation difference scores, or chemotaxis indices from all plates were pooled and metrics were compared across strains with ANOVA and Tukey honestly significant difference (HSD) tests. For all statistical tests an alpha value of 0.05 was used to determine significance.

## Acknowledgements

We would like to thank Dr. Evan L. Ardiel useful comments and discussion regarding the manuscript. We would like to thank Dr. Kurt Haas for motivating discussions to write the manuscript. We would like to thank Erica Li-Leger and the Moerman lab for assistance with experiments. We would like to thank Dr. John Calarco, Dr. Erik Jorgensen, and Dr. Hidehito Kuroyanagi and their labs for sharing their constructs and protocols or making them publicly available. We would like to thank Warren M. Meyers and Christine R. Ackerley for useful advice and discussions regarding figure design and/or protein structural modeling. We would also like to thank the *Caenorhabditis* Genetic Center for strains.

## Data availability

The data sets generated during the current study are available from the corresponding author on reasonable request. The code used to analyze data in the current study is available from the first author on reasonable request. All strains and reagents are available from the corresponding author upon reasonable request.

## Author Contribution Statement

TAM, VA, KM, and DGM conceived the genome editing strategy, designed and built constructs. TAM, VA, and JL generated the transgenic lines. TAM, VA, and KM performed genotyping and RT-PCR experiments. TAM and CHR designed the behavioral experiments. TAM, and ADL performed the behavioral experiments. TAM wrote custom scripts in R to organize the data and TAM and CHR analyzed the data. TAM wrote the first draft of the manuscript and made the figures. TAM, VA, KM, DGM, and CHR edited and co-wrote the final manuscript.

---

**Figure S1.**
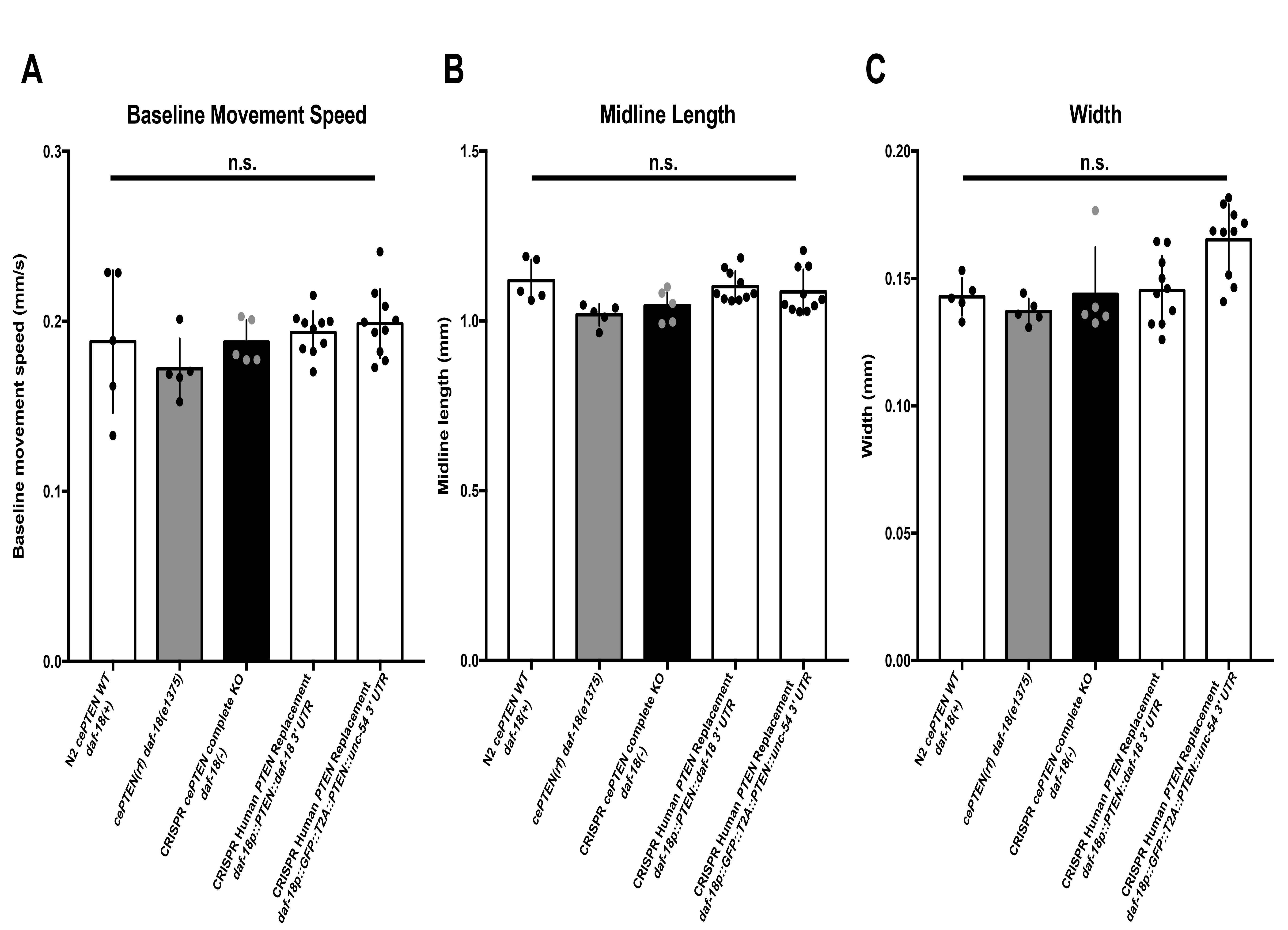
Morphology and baseline locomotion are superficially normal in *daf-18* mutants and *PTEN* transgenic animals. **A)** Baseline movement speed, **B)** midline length, and **C)** width are not significantly different across genotypes. Circles represent plate replicates run on the same day. Error bars represent standard deviation using the number of plates as n (n = 5 or 10). n.s. not significant, one-way ANOVA and Tukey’s post-hoc test.

